# Anti-inflammatory and neuroprotective activity of Viphyllin a standardized extract of β-caryophyllene from black pepper (Piper Nigrum L) and its associated mechanisms in mouse macrophage cells and Human Neuroblastoma SH-SY5Y cells

**DOI:** 10.1101/2021.12.16.472916

**Authors:** Kuluvar Gouthamchandra, Sudeep Heggar Venkataramana, Anusha Sathish, Amritharaj, Lingaraju Harakanahalli Basavegowda, Naveen Puttaswamy, Shyam Prasad Kodimule

## Abstract

Oxidative stress breeds various chronic lifestyle ailments including inflammatory conditions and neurodegenerative diseases. β-caryophyllene natural bicyclic sesquiterpene, obtained from various plants sources found to be effective against inflammation and neuroprotection. In this study, we have evaluated the protective effect of Viphyllin, a standardized extract of β-caryophyllene from black pepper against inflammation induced by lipopolysaccharide in RAW264.7 macrophage cells and mechanisms involved in hydrogen peroxide (H_2_O_2_)-challenged oxidative stress in human neuroblastoma SH-SY5Y cells. Viphyllin demonstrated the anti-inflammatory activity by subsiding the release of the pro-inflammatory intermediaries like NO, cytokines, interleukins, and protein expression levels of cyclooxygenase (COX-2) and inducible nitric oxide synthase (iNOS). In addition, Viphyllin suppressed the extracellular signal-regulated kinase (ERK). c-Jun N-terminal kinase (JNK) and p38 mitogen-activated protein kinase (MAPK) phosphorylation. On the other hand, Viphyllin showed neuroprotective effect against neuronal oxidative damage caused by H_2_O_2._ Viphyllin lessened the expression B-cell lymphoma-2 (Bcl-2), B-cell lymphoma-2-associated X protein (BAX), cleaved caspase-9, and PARP-1 proteins associated with apoptosis. Our results indicate that Viphyllin ameliorated LPS-mediated inflammation in macrophages by regulating inflammation and Viphyllin exerted remarkable anti apoptotic effect against neuronal damage challenged by H_2_O_2_. Altogether, Viphyllin could be potential functional food ingredient for inflammation and neurodegenerative diseases.

## Introduction

Pharmacological properties of medicinal plants are primarily due to the presence of secondary metabolites like polyphenols, polyphenols, steroids, flavonoids, alkaloids and terpenoids [1,2]. Terpenoids have been studied as an alternative to treat different disease conditions. Certain terpenoids for instance Taxol and Artemisinin has been using as anticancer and antimalarial drug, respectively. Terpenoids play diverse role in the field of food, drugs, and cosmetics, studies show the relationship of terpenoid consumption with the reduction of inflammation [3,4]. β-caryophyllene is a “natural bicyclic sesquiterpene hydrocarbon” it is most abundant constituent exist in many plant-originated essential oils, like as lemon balm, cloves, Cannabis sativa and black pepper [5,6]. Recent studies are reported on significant biological activities of β-caryophyllene against inflammation, cancer and chronic pain and chemo sensitizing properties for chemotherapies like oxaliplatin, doxorubicin sorafenib, and 5-fluorouracil. On the other hand, β-caryophyllene is a plant (or food)-derived cannabinoid ligand, and it is also reported that it functions as a selective agonist of cannabinoid receptor 2 (CB2). It is thought that CB (2) receptor activation is suitable medicinal approach for the treatment of inflammation related disorders. United States Food and Drug Administration considers β-caryophyllene is a dietary phyto cannabinoids and has been included in the list of “generally regarded as safe” ingredients for dietary use and [7,8].

In the current investigation, we have enriched and systematized the extract in a manner that is not less than 30% β-caryophyllene and termed it as Viphyllin, subsequently investigated for anti-inflammatory activity in LPS induced mouse macrophage cells and neuroprotective property against oxidative stress induced by H_2_O_2_. To the best of our knowledge, this is the very first study revealing that a standardized extract of beta caryophyllene (Viphyllin) from black pepper demonstrated an anti-inflammatory and neuroprotective activity.

## Materials and methods

### Sample preparation

Viphyllin a standardized extract of β-caryophyllene from black pepper (Piper Nigrum L) was collected from the Department of Phytochemistry, Vidya Herbs Pvt. Ltd. (Bangalore, Karnataka, India), and dissolved in cell culture media (DMEM) at the appropriate concentrations. (**Supplementary figure S1**)

### Cell culture

RAW 264.7 Murine macrophage and Human neuroblastoma SH-SY5Y cells were obtained through NCCS (Pune, India), RAW264.7 cells were cultured in DMEM media containing 10% foetal bovine serum and antibiotic solution in a humidified 5% CO_2_ incubator (Thermo Fisher Scientific). The cell culture medium was replaced with fresh DMEM media every 2 to 3 days, once cells reached 80% confluence, the cells were subcultured. SH-SY5Y cells were grown in DMEM and F12 nutrient mix subsidized with FBS (10%) and antibiotic solution respectively at 37 °C in the CO_2_ incubator.

### Cell viability assay

MTT assay was adopted to check the influence of Viphyllin on the viability of RAW 264.7 cells. Briefly, cell suspensions were seeded at density of 1×10^4^ cells/well in 96-well plates. After 18 h of incubation, the cells were treated with the different concentrations(5-50μg/mL) of Viphyllin, further incubated for 24 h. The supernatant was discarded and 100 μL of MTT solution (0.5 mg/mL) was replaced and cells were incubated for an extra 2-4hrs, subsequently 100 μL of MTT stop solution (DMSO) was added to wells to dissolve the formazan crystal. The absorbance was read at 550 nm using a microplate reader (Tecan Infinite M200 Pro). Likewise, cell viability assay was conducted on SHSY5Y cells. The preventive effect of Viphyllin versus H_2_O_2_-elicited neurotoxicity was investigated by incubating cells with 12 and 24µg/mL of Viphyllin for 6 h and then treated them with 500μM H_2_O_2_ for 24 h. Cell viability of cells were determined using the MTT assay as described above.

### Determination of NO

RAW macrophages were set at 4×10^4^ cells/well in 96-well plates for 18 h. Then the cells were incubated with different concentrations (5-50μg/mL) of Viphyllin before they were stimulated with 1μg/ml lipopolysaccharide. After completion of 24h incubation, using the cell culture supernatant NO content was measured by Griess reagent.

### Evaluation of cytokine secretion level in macrophages

The secretion level of PGE-2, TNF-α, IL-6 and IL-1β, inflammatory cytokines levels were measured by the ELISA using in the cell culture supernatant of RAW 264.7 macrophages according to manufacturer’s protocol (Krishgen Biosystems, Mumbai, India).

### Lactate dehydrogenase (LDH) assay

The assay was performed by employing an LDH cytotoxicity detection kit (Sigma aldrich). SH-SY5Y cells were added at the density of 2.5×10^4^ cells/ mL in 96-well. Then the cells were treated with Viphyllin at different concentrations (5-50μg/mL) for 24h. with and without H_2_O_2_. The amount of LDH release was determined by using a microplate reader (Tecan Infinite M200 Pro) at 490 nm.

### Western Blot Analysis

RAW macrophages were incubated overnight and subjected to two different concentrations of Viphyllin 12 and 24µg/mL prior to LPS (1µg/mL) treatment. Cells were collected by trypsinization after 24 h incubation and were lysed with 100µL of radioimmunoprecipitation (RIPA) lysis buffer. Total protein concentration was measured using the Bradford reagent (Bio-Rad). Equivalent amounts of protein (30µg) were separated by sulfate-polyacrylamide gel electrophoresis (SDS-PAGE), and the protein bands were transferred onto a polyvinylidene fluoride (PVDF) membrane. The membrane was blocked with 5% non-fat milk solution (and incubated with 1:1000 diluted primary antibodies (cox2. iNOS, p-ERK, p-JNK, p-p38 and β-actin) at 4°C for overnight, and subsequently probed with specific secondary antibodies coupled to horseradish peroxidase (Santa Cruz Biotech, Dallas, TX, USA) and bands were visualized using ECL reagents, and quantified using the ImageJ software (National Institutes of Health, Bethesda, MD, USA). Similarly, western blot experiment was conducted in H_2_O_2_ induced oxidative damage in SH-SY5Y cells using PARP1, caspase 9, Bcl-2 and BAX primary antibodies.

### Statistical analysis

One-way ANOVA and Dunnett’s multiple comparisons test has been used to determine the statistical significance of differences among control and treated groups. Experiments were carried out in triplicate and all the values that were expressed as a mean ± standard error.

## Results

### Cell viability

We investigated the cytotoxic effect of Viphyllin in RAW264.7 murine macrophages. As shown in Figure 1A. Viphyllin had no cytotoxic effect on macrophages up to the dosage of 30 µg, whereas higher dose of Viphyllin showed significant cytotoxicity in RAW264.7 cells. Hence, we concluded that concentrations up to 30 μg/ml of the Viphyllin could be safe. Thus, in the subsequent experiments, RAW274.7 cells were treated with nontoxic concentrations of Viphyllin. The morphology of LPS elicited cells had an uneven shape and spreading compared to untreated cells, whereas Viphyllin treated cells offer protection against LPS treatment by maintaining normal shape of cells (Figure 1B).

**Figure 1.**
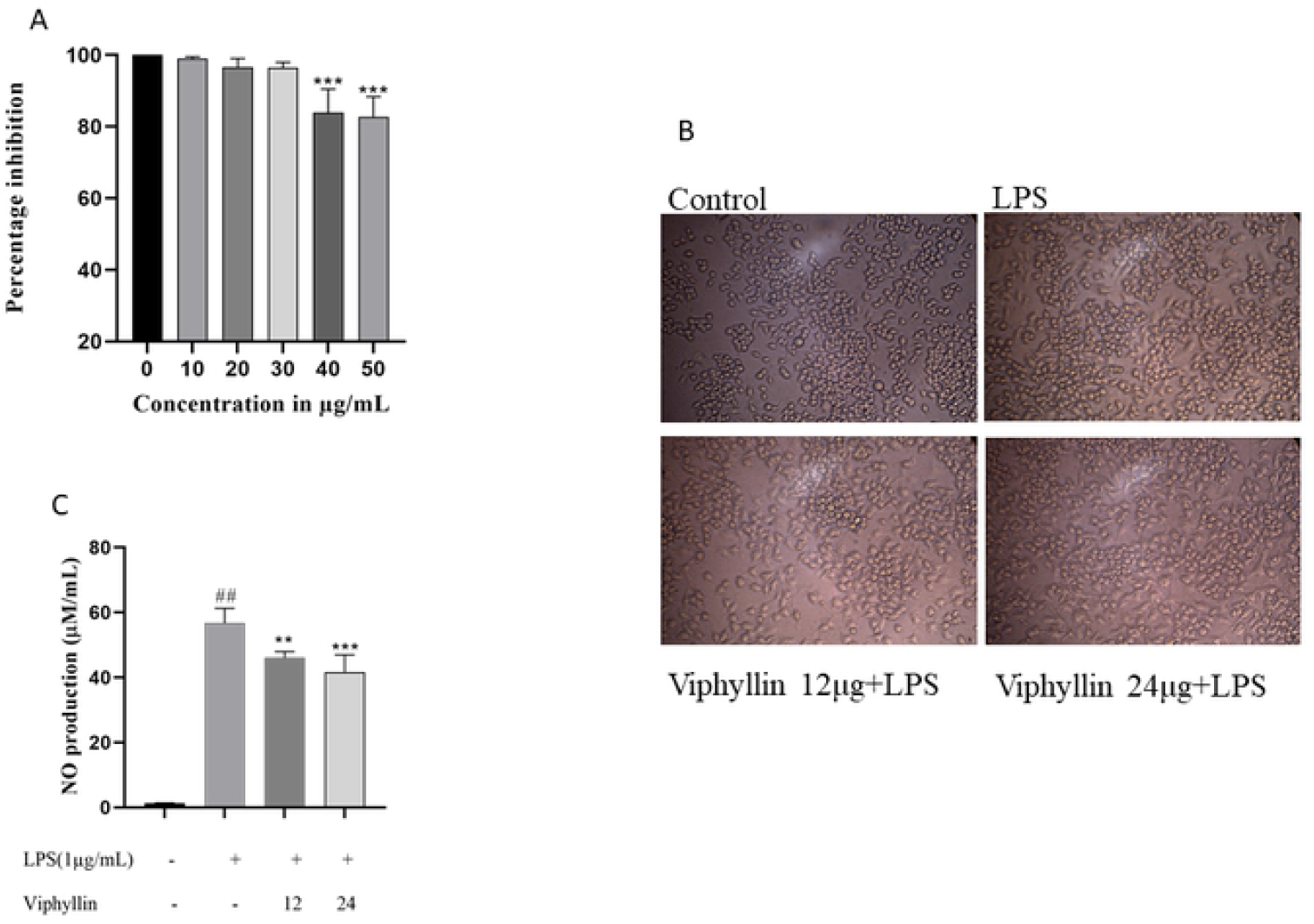

### Effect of Viphyllin on Nitric Oxide (NO) production

NO is secondary intermediate secreted by macrophages upon LPS stimulation which plays significant role in regulating natural and genetic immunity. Here in we observed the repressive activity of Viphyllin on production NO, the levels of NO production in the cell culture supernatant of LPS provoked RAW 264.4 murine macrophages after 18 h exposure were high and observed that Viphyllin treatment significantly decreased production of NO levels in macrophages when compared to LPS (positive control) treated cells. (Figure 1C).

### Influence of Viphyllin on secretion of PGE2, TNF-α. IL-6 and IL-1β in LPS challenged RAW264.7 macrophages

The effect of Viphyllin treatment on the secretion of inflammatory mediators PGE2 and cytokines (TNF-α IL-6 and IL-1β) in the supernatant of RAW264.7 macrophages challenged with LPS were examined. As shown in Figure 2A-D, LPS significantly elevated secretion level of cytokines in macrophages *(P*-value<0.001 respectively) compared normal cells. Whereas Viphyllin prior treatment significantly diminished the LPS elicited production of proinflammatory cytokines TNF-α, PGE-2, IL-6 in a concentration reliant manner, whereas IL-1β expression remains unchanged.

**Figure 2.**
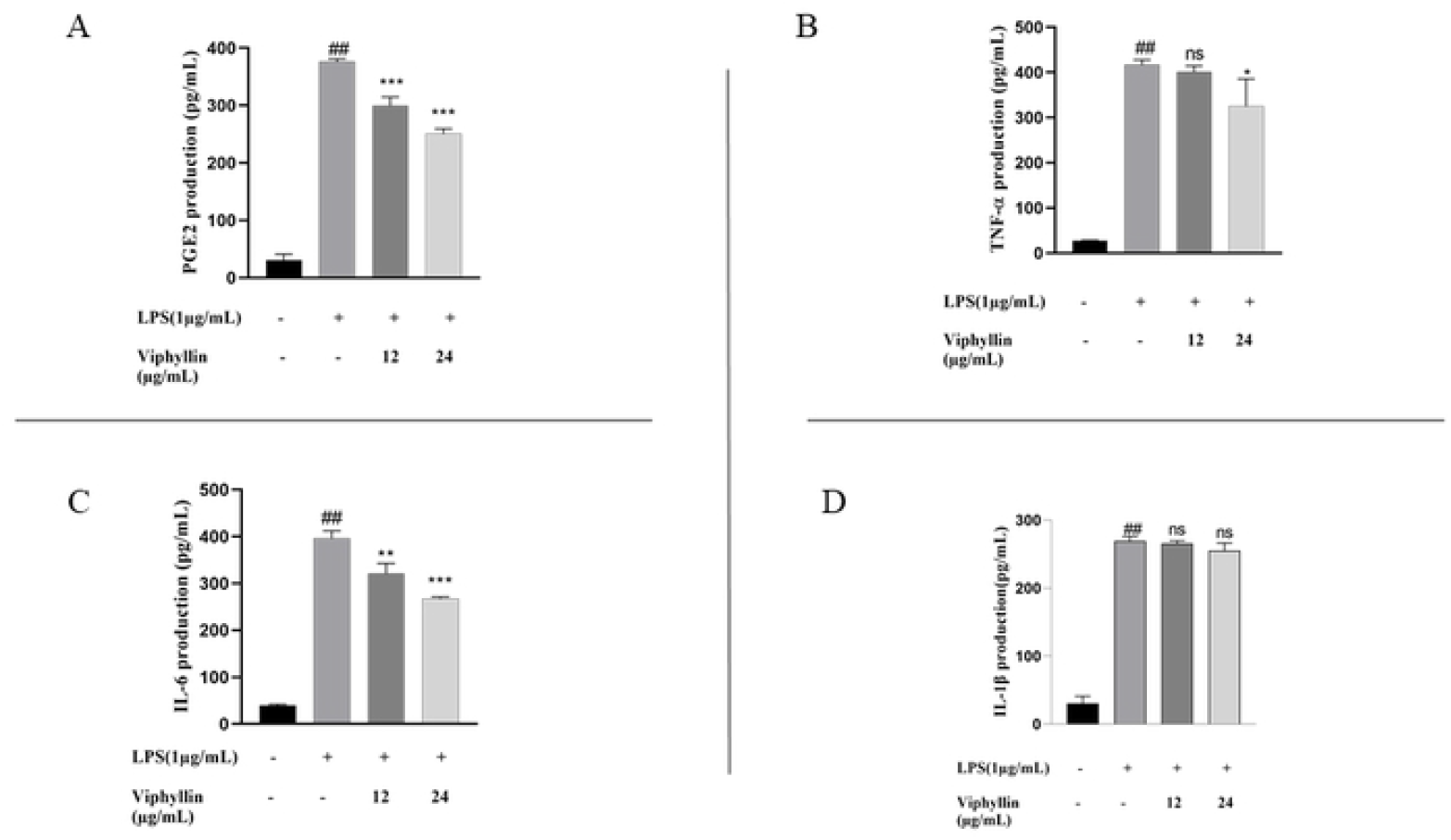

### Effect of Viphyllin on the expression amount of COX-2, and iNOS protein in LPS elicited RAW 264.7 macrophages

To examine the influence of Viphyllin as anti-inflammatory agent we have carried out western blotting experiments to check the expression amounts of COX-2, and iNOS protein in murine macrophages challenged by LPS(1µg/ml). 24h later, the presence of LPS (1µg/ml) in RAW 264.7 cells caused the dramatic upregulation of COX-2 and INOS protein levels when compared untreated normal cells. However, treatment with Viphyllin, in presence of LPS significantly subsided the expression levels of COX-2 and iNOS proteins at concentration at 24µg/mL compared to LPS alone treated RAW 264.7 macrophages (Figure 3).

**Figure 3.**
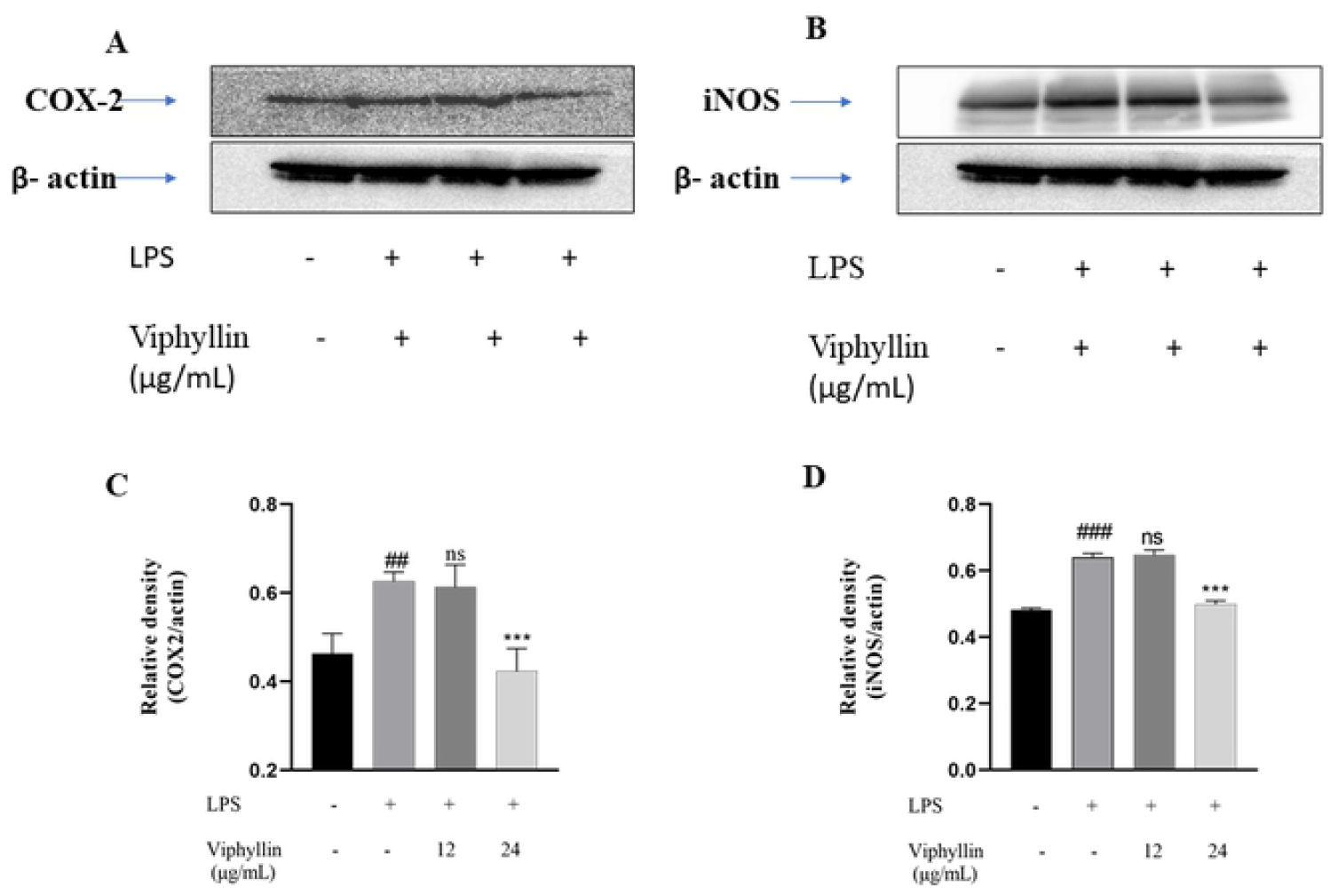

### Effect of Viphyllin on MAPK signalling pathway in LPS-Induced RAW 264.7 Cells

In many inflammatory diseases including cancer, viral infection and neurological disorders, extreme inflammation is pretty established as a crucial factor. Enhanced activity of MAPK, specifically p38 and their involvement in the management of the synthesis of inflammation intermediaries make them targets for anti-inflammatory drugs. To examine if these signalling pathways are affected by Viphyllin, we conducted western blot experiments. Western blot results showed that LPS stimulation significantly elicited phosphorylation of p-ERK, p-JNK and p-p38 MAPK compared to control untreated cells. In contrast we noticed Viphyllin pre-treatment mitigated phosphorylation of JNK and p38 (Fig. 4A-D). However, interestingly we observed Viphyllin at the concentration of 12µg/mL had significant effect on suppression of p-ERK. These results suggest that Viphyllin could inhibit inflammatory response through regulation of MAPKs signalling in LPS-activated murine macrophages.

**Figure 4.**
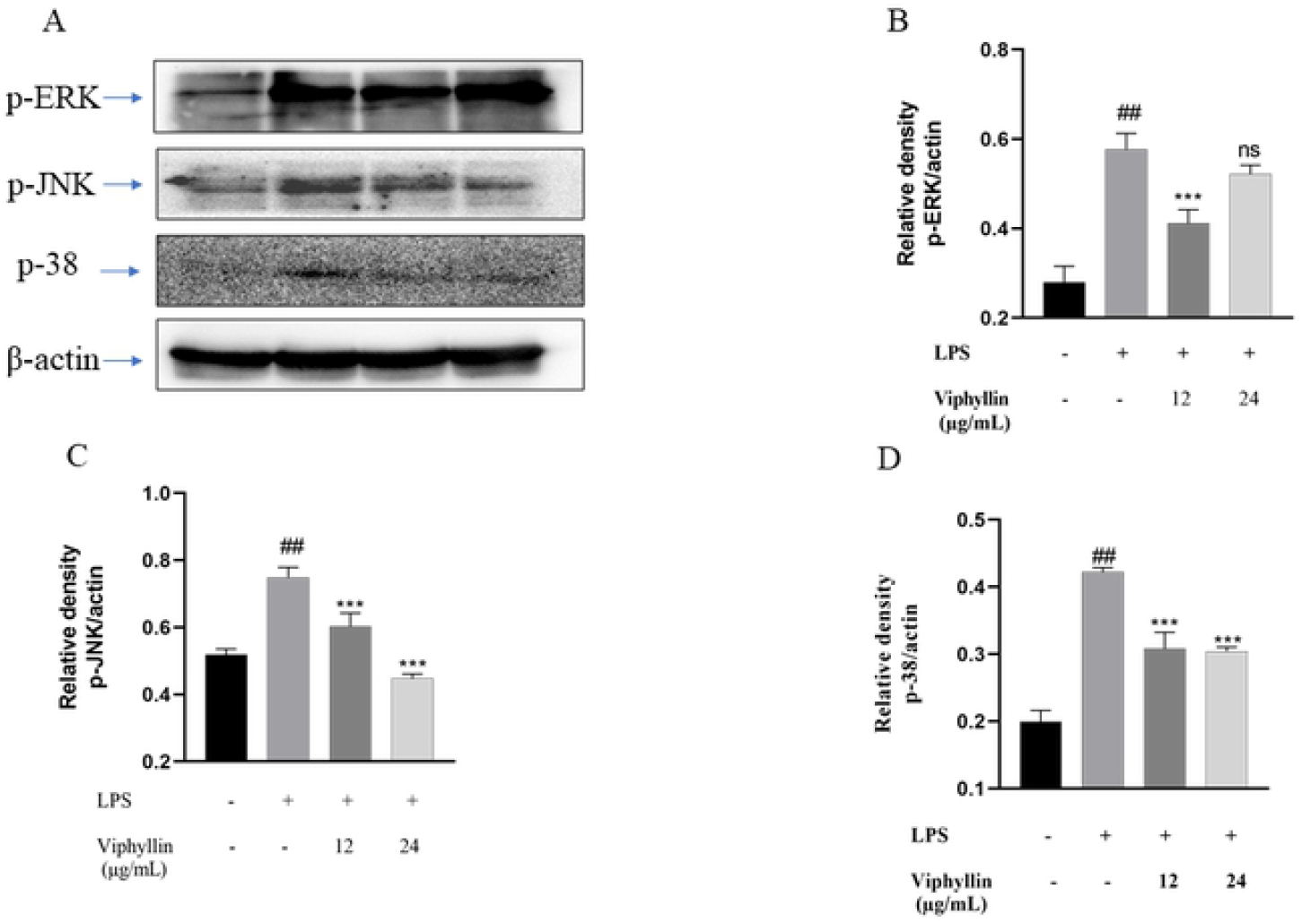

### Neuroprotective Effect of Viphyllin on H_2_O_2_-challenged Human Neuroblastoma SH-SY5Y cells

Investigate the effect of Viphyllin presence on cell viability, SH-SY5Y cells were treated with Viphyllin at various concentrations ranging from 5–50µg/mL for 24 hr (Figure 5A). It has been observed that Viphyllin had no significant effect on SH-SY5Y cells up to the dosage of 30μg/ml where its viability ranges from (98.3, 95.97% and 95.12%) respectively, whereas higher concentration of Viphyllin subsided the cell viability. Therefore, we have selected the nontoxic dose (12 and 24μg/ml) of Viphyllin for further experiments. H_2_O_2_ is well established neurotoxin, which generates highly reactive oxygen species like hydroxyl, superoxide anion radicals, resulting in cellular damage. Determine protective effect of Viphyllin against H_2_O_2_ induced oxidative stress in SH-SY5Y cells, cells were pre-incubated with two different concentration (12 and 24μg/mL) of Viphyllin for 4 h and then subjected to 500μM of H_2_O_2_ for 24 h. cell viability assay demonstrated that presence of H_2_O_2_ decreased viability of SH-SY5Y cells, while interestingly 24μg/mL of Viphyllin pre-treatment showed considerable protection against H_2_O_2_-generated loss of cells (Figure 5B).

**Figure 5.**
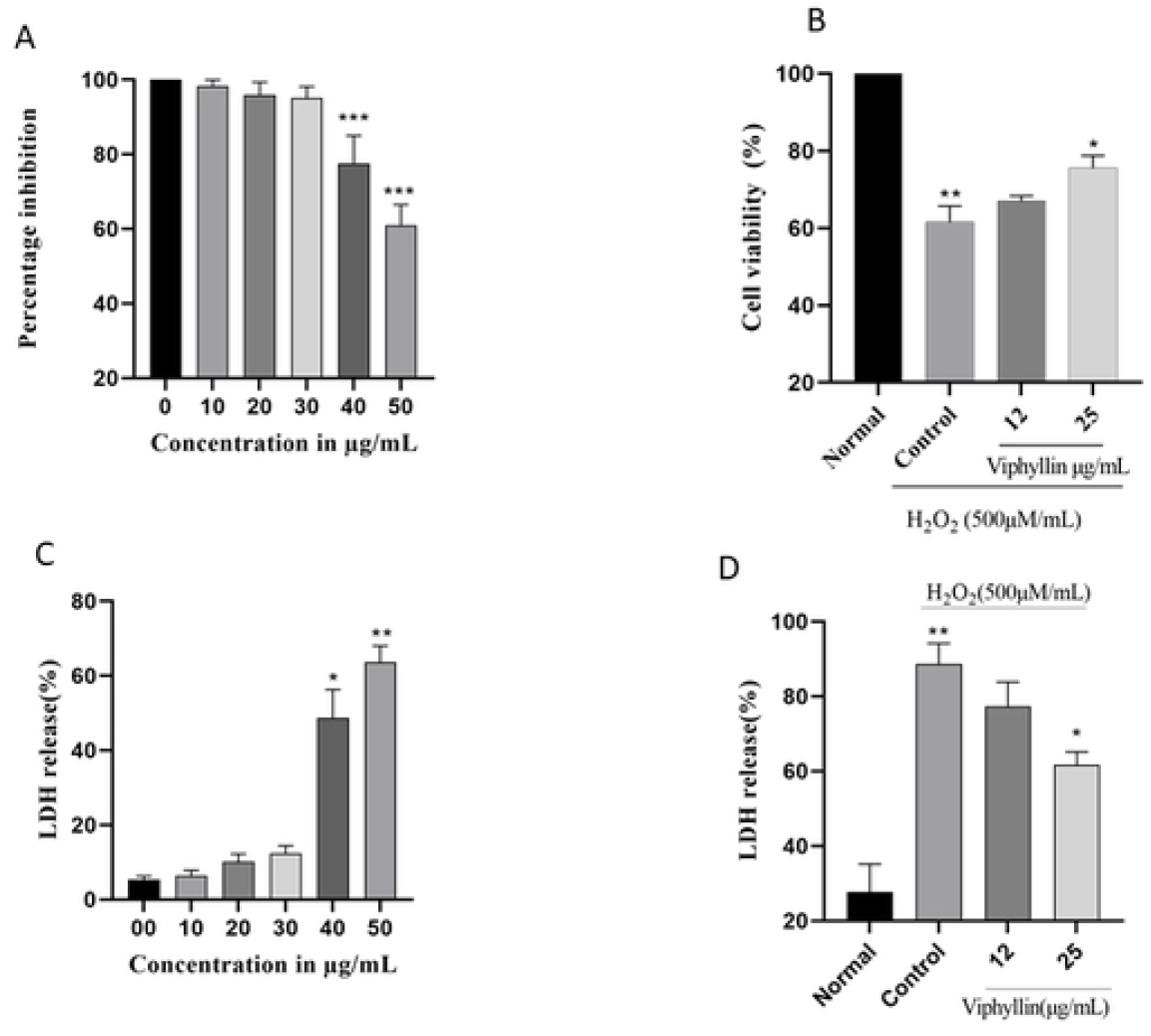

### LDH assay

LDH is an intracellular enzyme that exists in the cells, elevated levels of LDH enzyme in the cell culture medium implies cellular membrane burst and it is considered as label of cell death. Impact of various concentration of Viphyllin on LDH release was examined, as shown in Figure 5C. Viphyllin up to the concentration of 30μg/mL did not affect the LDH release, on higher concentration significant release of LDH was observed, consequently nontoxic dose of Viphyllin was evaluated for its protective effect against H_2_O_2_ induced toxicity. LDH secretion in the H_2_O_2_-treated control group was significantly elevated compared to untreated control cells with the release of 88.66%. However, pre-treatment of Viphyllin significantly decreased H_2_O_2_-induced LDH release suggesting suppression of neuronal toxicity stimulated by H_2_O_2_ in SH-SY5Y cells (Figure 5D).

### Effects of Viphyllin on apoptosis-related protein expressions in H_2_O_2_-induced SH-SY5Y cells

It has been reported that accumulation of H_2_O_2_ causes apoptosis, inflammation and necrosis in the brain is well known in neurodegenerative disease. H_2_O_2_ stimulated oxidative stress in SHSY-5Y cells is well established model for studying neurological disorders. Determine the antiapoptotic effect of Viphyllin, we investigated the of pro and antiapoptotic protein expression in SHSY-5Y cells. As presented in Figure 6A-C. we found the increased expressions of cleaved PARP1 and caspase-9 in the H_2_O_2-_stimulated cells when compared with the untreated normal cells. However, in the treated cells groups, Viphyllin lessened the - highly expressed cleaved PARP1 and caspase-9 in a concentration reliant manner. In addition, As represented in Figure 6 D-F. H_2_O_2_(500µM) treated group we found significantly elevated expression of the pro-apoptotic gene BAX and subsided expression of Bcl-2 an anti-apoptotic protein in SH-SY5Y cells, whereas Viphyllin treatment dose dependently reversed these effects.

**Figure 6.**
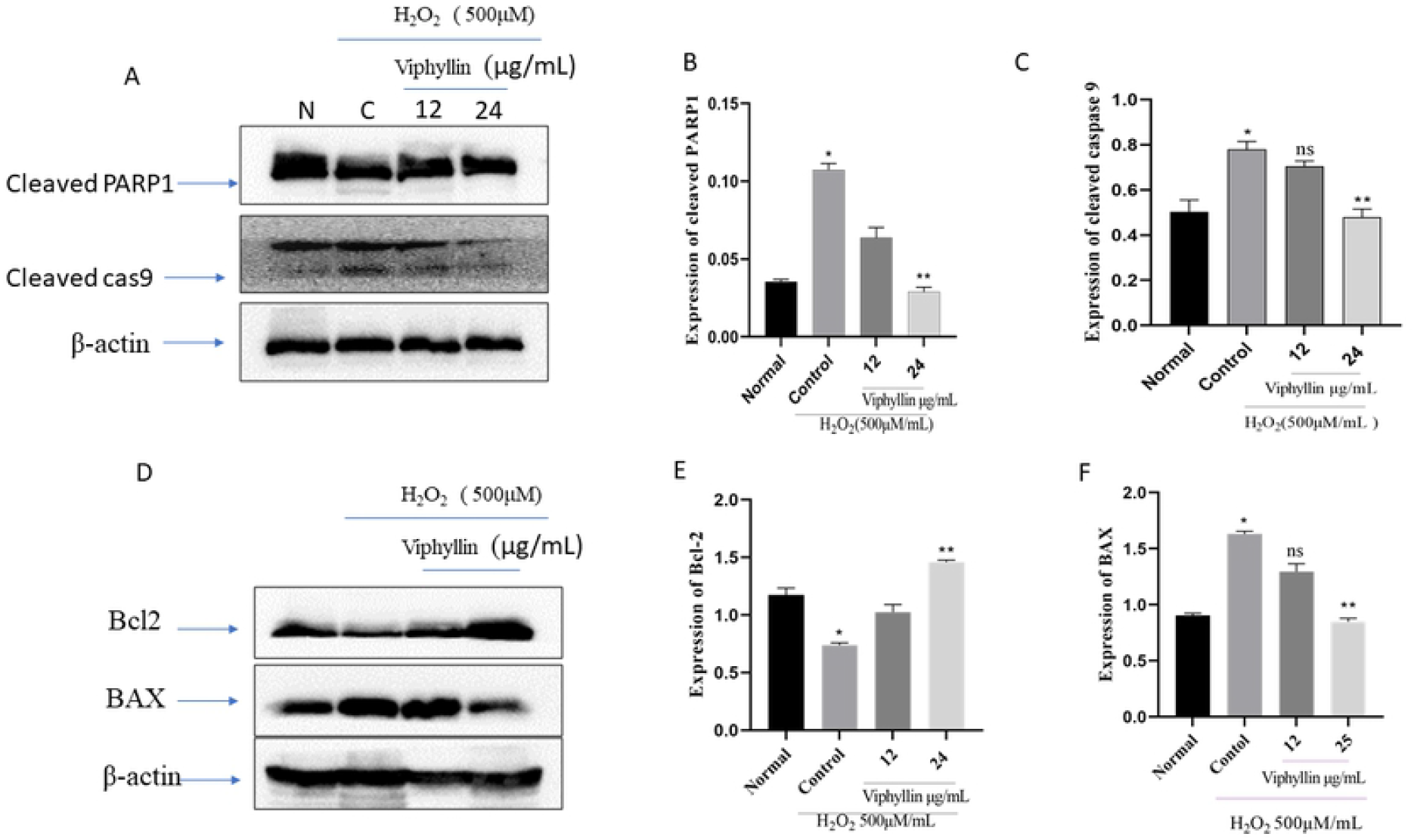

## Discussion

Persistent inflammation has a extensive role in the pathogenesis of various chronic diseases like diabetes, obesity, cardiovascular and neurological disorders [9–11]. Approaches to aim for inflammation has become an attractive approach to defend inflammatory-related ailments [12,13]. Over the last few years, bioactive constituents or standardized extracts obtained from plant sources still play a significant role in the treatment of many ailments without any adverse effect [14]. Macrophages are crucial participants in the immune response to foreign intruders such as infectious microorganisms. lipopolysaccharide (LPS) is a well-known prototypical endotoxin may directly activate macrophages. Plethora of measures have recommended that activated macrophage produce a greater amount of various proinflammatory intermediaries like NO and PGE2, also proinflammatory cytokines for example TNF-α, and interleukins. All these elements stimulate the development of inflammation and worsen the inflammation over the harmonious interaction with further inflammatory intermediaries and there are reports that in variety of inflammatory ailments including asthma and colitis the expression levels of PGE2 and NO is elevated [15–17]. Our study revealed that Viphyllin efficiently inhibited the secretion of PGE2 and NO, and proinflammatory cytokines in LPS-challenged macrophages at safer dose, our findings are consistent with earlier published studies [18, 19]. COX-2. is an important target for the design of drugs against inflammation, particularly NSAIDs. Furthermore, iNOS and COX-2 enzymes play crucial role in the generation of NO and PGE2, Viphyllin significantly subsided the iNOS and COX-2 expression in LPS stimulated macrophages. Our results are consistent with the previous published reports [20,21].

The MAPK signalling cascade plays a key role in the initiation of response to inflammation. ERK, JNK, and p38 participate in MAPK signalling pathways, which gets activated by TNF-α and interleukins [22]. Earlier studies have shown that LPS or TNF activates MAPKs which are involved in acute lung injury. It has been reported that when MAPK gets activated phosphorylated ERK, p38, and JNK are moved to the nucleus and stimulates response to inflammation by inducing expression of proper gene of interest [23,24]. Pre-treatment of Viphyllin inhibited MAPK signalling by subsiding the phosphorylation of ERK, JNK, and p38 in LPS induced macrophages. It is evident from the study that inhibition of MAPK pathway is a potential mechanism by which Viphyllin wields anti-inflammatory effects. These findings are consistent with those earlier studies [25,26].

Lately, several findings have found neurodegenerative diseases caused by stress and has been extensively associated in neuronal cell death. Previous studies have reported that H_2_O_2_ accumulation has been well studied in neurodegenerative disease. Human neuroblastoma SH-SY5Y cells are very well-regarded cellular tool for research on H_2_O_2_-stimulated oxidative stress ^27–29^. In our recent study we have explored standardized β-caryophyllene extract improves cognitive function in scopolamine-induced amnesia model [30]. In current study, we have studied the protective effect of Viphyllin against H_2_O_2_-provoked apoptotic events in SH-SY5Y cells. It has been showed that H_2_O_2_ stimulated oxidative damage dramatically declines viability of SH-SY5Y cells [31]. Our study also showed decline of cell viability in the presence of H_2_O_2_ in SH-SY5Y cells. Interestingly we have noticed viability of cells enhanced dose dependently by the treatment of Viphyllin. It has been well established that damage to cellular membranes releases LDH into the cell culture supernatant is important marker of cell death [32]. From our data it is noticeably clear that elevated level of LDH was found in the culture medium after H_2_O_2_-induction in SH-SY5Y cells. Treatment with the Viphyllin markedly lower the LDH in the SH-SY5Y cell culture medium subjected to H_2_O_2_. Hence, we reaffirmed the defensive effects of Viphyllin in H_2_O_2_-elicited SH-SY5Y cells. Our outcomes are constant with previously published study [33]. It has been reported that toxic effect on SHSY-5Y cell is caused by H_2_O_2_ is linked with mitochondria-associated apoptotic events and it is considered as one of the valuable tools in neurodegenerative diseases [34]. Cytochrome release occurs when elevated levels of ROS penetrate mitochondrial membrane which leads to the activation of caspase-9 and PARP-1 and gradually induces cell death [35]. Our results demonstrated that markedly elevated proteolytic cleavage of caspase-9 and PARP in H_2_O_2_ subjected SHSY-5Y cells. However, Viphyllin protects neuronal cells against H_2_O_2_ stimulated oxidative stress by lessening the expression of caspase-9 and PARP1. Furthermore, mitochondrial membrane injury leads to neuronal apoptosis by up-regulation of Bcl-2 and down-regulation of Bax, which leads to inhibition of caspase activation [36]. In current study, Viphyllin treatment lowered Bax level and enhanced Bcl-2 level. Hence, these outcomes demonstrated that Viphyllin along with other active constituents from black pepper offered preventive effects against oxidative damage, our results were consistent with other published studies [37,38]. In conclusion, our current unearthing demonstrated that the Viphyllin standardized extract containing β-caryophyllene from black pepper exert an anti-inflammatory function via the downregulation COX-2, iNOS and of the MAPK pathway in LPS challenged macrophages on the other hand it also offers neuroprotective effect. This investigation may suggest Viphyllin is encouraging bioactive food ingredients. However, imminent research is essential relating to the other controlling mechanism of Viphyllin.

## Acknowledgements

We would like to thank, Mr. Chandrappa S, Dr. Deepak, and Dr. Vedamuthy B.M, Department of Phytochemistry and Analytical development laboratory, Vidya Herbs Pvt. Ltd, Bangalore, India. Our lab members are also acknowledged for their useful suggestions.

## Authors’ contributions

GK and SK drafted the original manuscript. GK, SHV, AS and AR Produced the data and prepared figures 1–6. LH and N P prepared figure S1. GK, SK and SHV designed and analyzed the experiment and results. All authors read and approved the final manuscript.

Quan L, Thiele G, Thian J, Wang D (2008) The development of novel therapies for rheumatoid arthritis. Expert Opin Ther Pat. 18: 723–738

